# Cycling in degradation of organic polymers and uptake of nutrients by a litter-degrading fungus

**DOI:** 10.1101/2020.07.07.170282

**Authors:** Aurin M. Vos, Robert-Jan Bleichrodt, Koen C. Herman, Robin A. Ohm, Karin Scholtmeijer, Heike Schmitt, Luis G. Lugones, Han A. B. Wösten

**Affiliations:** Microbiology, Department of Biology, Utrecht University, Utrecht, The Netherlands; Wageningen Plant Research, Wageningen UR, Wageningen, the Netherlands; Plant Breeding, Wageningen University and Research, Wageningen, The Netherlands; Institute for Risk Assessment Sciences, Utrecht University, Utrecht, The Netherlands

**Keywords:** fungi, Agaricus bisporus, litter degradation, lignocellulose, respiratory burst, cycling

## Abstract

Wood and litter degrading fungi are the main decomposers of lignocellulose and thus play a key role in carbon cycling in nature. Here we provide evidence for a novel lignocellulose degradation strategy employed by the litter degrading fungus *Agaricus bisporus* (known as the white button mushroom). Fusion of hyphae allows this fungus to synchronize the activity of its mycelium over large distances (50 cm). The synchronized activity has an 13-hour interval that increases to 20 h before becoming irregular and is associated with a 3.5-fold increase in respiration while compost temperature increases up to 2 °C. Transcriptomic analysis of this burst-like phenomenon supports a cyclic degradation of lignin, deconstruction of (hemi-) cellulose and microbial cell wall polymers, and uptake of degradation products during vegetative growth of *A. bisporus*. Cycling in expression of the ligninolytic system, enzymes involved in saccharification, and nutrient uptake is proposed to provide an efficient way for degradation of substrates such as litter.

## Introduction

Mushroom forming fungi play a key role in nature and human society(Grimm and Wösten, 2018). For instance, they produce edible and medicinal fruiting bodies and play a pivotal role in cycling of carbon in nature. The latter is illustrated by the fact that they are the main decomposers of lignocellulose, the most abundant terrestrial organic material. This complex is found in wood and litter and consists of cellulose, hemicellulose and lignin. Brown rot fungi mostly modify lignin and degrade carbohydrates while white rots simultaneously degrade lignin and carbohydrates or selectively (i.e. predominantly) degrade lignin (Worrall *et al*., 1997; Riley *et al*., 2014; Schilling *et al*., 2015). Brown and white rot wood degrading fungi use different types of radical chemistry to modify and degrade lignin (ten Have and Teunissen, 2001; Hammel *et al*., 2015), thereby exposing (hemi-) cellulose to the action of carbohydrate-active enzymes (CAZYs).

Recently, it was found that the mycelium of mushroom forming brown rot fungi spatially separate the radical generating Fenton chemistry from CAZYs (Zhang *et al*., 2016, 2019; Presley and Schilling, 2017; Castaño *et al*., 2018). The outer 5 mm of the colony of brown rots like *Postia placenta* generate •OH radicals using the Fenton reaction, while the inner 15-35 mm regions of the colony secrete the bulk of (hemi-) cellulases. This spatial separation would prevent inactivation of these enzymes by radical oxygen species generated by Fenton chemistry and the radical pre-treatment of cell walls may facilitate the activity of CAZYs (Castaño *et al*., 2018; Zhang *et al*., 2019).

White rots may also make use of Fenton chemistry to attack lignocellulose but it is not the main mechanism. In this fungal decay type lignin is mineralized using a ligninolytic system that includes oxidoreductases like lignin peroxidase, manganese peroxidase (MnP), and / or versatile peroxidase, and possibly laccase (LCC) as well (ten Have and Teunissen, 2001; Hammel *et al*., 2015). The peroxidases use H_2_O_2_ that is generated by oxidases like glyoxal oxidase but it may also arise from secondary reactions with organic acids like oxalate (Kersten and Kirk, 1987; Urzúa *et al*., 1998). As a result, small radical intermediates are produced that penetrate lignocellulose and attack lignin directly or indirectly by facilitating secondary reactions, possibly including lipid peroxidation in the case of MnP (Hofrichter, 2002). Radicals generated by the ligninolytic system of white rots may result in a more gentle radical treatment of lignocellulose as not all radicals species that are associated with white rot match the reactivity of •OH radicals. This may explain why white rots have been described to produce their ligninolytic system together with (hemi-) cellulases. For example, the white rot *Trametes versicolor* expresses genes related to ligninolytic and (hemi)-cellulolytic activity in a region 15-35 mm from the colony hyphal front (Presley *et al*., 2018; Zhang *et al*., 2019). The white rots *Phanerochaete chrysosporium* and *Stereum hirsutum* have a delayed onset of ligninolytic activities (Moukha *et al*., 1993; Korripally *et al*., 2015; Presley *et al*., 2018) and expression of (hemi-) cellulolytic genes in *P. chrysosporium* occurs in both younger and older mycelium (Korripally *et al*., 2015). In contrast, *Pleurotus ostreatus* separates the bulk of (hemi-) cellulases from its ligninolytic system by producing (hemi-) cellulases 15-20 mm from its hyphal front while its ligninolytic system is most active in the 0-5 mm and 30-35 mm region of its colony (Zhang *et al*., 2019). Together, white rots show differences in their spatial expression of ligninolytic and (hemi)-cellulolytic activities.

Soil-inhabiting saprotrophs like the litter-decomposer *Agaricus bisporus* (the white button mushroom) are, like white rots, capable of extensive lignin degradation (Osono, 2007; Jurak *et al*., 2015). The genomes of litter-degrading and white rot fungi indicate differences in their ligninolytic machinery, possibly relating to the difference in wood and litter environments (Floudas *et al*., 2020). Yet, the enzymatic system enabling the decomposition of lignin by *A. bisporus* includes MnP and LCC that are thought to play an important role in lignin breakdown by some white rot fungi as well. The expression of CAZYs and their activities throughout the life cycle of *Agaricus bisporus* have been relatively well studied. This fungus is grown commercially on a horse manure based compost (Grimm and Wösten, 2018) that is produced in two phases. *A. bisporus* is introduced at the end of Phase II (PII), which starts the vegetative growth period during Phase III (PIII). After topping the PIII-end compost with a casing layer and changing environmental conditions, fruiting is induced (Phase IV [PIV]). It is generally assumed that *A. bisporus* degrades the lignocellulose in the substrate by means of (hemi-) cellulases, LCC and MnP (Bonnen *et al*., 1994; Patyshakuliyeva *et al*., 2015). About 40 % of the lignin, 6 % xylan (part of the hemicellulose), 6 % cellulose, and a significant part of the microbial population that develops during the preparation of PII compost is consumed during PIII relative to PII (Kabel *et al*., 2017; Vos, Heijboer, *et al*., 2017). Xylan and cellulose are preferentially degraded during PIV with approximately 48 % and 33 % cellulose and xylan being removed after fruiting, respectively, while minor amounts of lignin are removed. Loss of lignocellulose in compost in time is in agreement with overall expression patterns of ligninolytic and (hemi-) cellulolytic genes (Patyshakuliyeva *et al*., 2015).

So far, physiology and gene expression studies have not been performed at short time intervals during the different phases of the life cycle of *A. bisporus* such as during PIII. Here, we performed such studies, revealing that the expression profile of *A. bisporus* during its vegetative growth is in line with its mycelium cycling through degradation of lignin, degradation of polysaccharides, and uptake of degradation products. This cycling, which is accompanied by respiratory bursts, is synchronized over a large distance within the mycelium and is proposed to enable *A. bisporus* to efficiently degrade litter.

## Material and Methods

### Strain and growth conditions

*A. bisporus* strains A15 (Sylvan, Netherlands) and Bisp015 were grown in PII-end compost (CNC Grondstoffen, Milsbeek, the Netherlands) at 25 °C unless otherwise indicated. Compost was inoculated with either rye based A15 (Gift from CNC Grondstoffen, Milsbeek, the Netherlands) or Bisp015 spawn (Basidiomycete culture collection WUR, Wageningen, the Netherlands) or with PIII compost. Respiration measurements were done in 1 litre bottles using 50 g PII-end compost that was either or not inoculated with 10 spawn grains. Temperature was monitored either in boxes or in square Petri dishes. In the latter case, square Petri dishes (12 × 12 × 1.2) were filled with 57.6 g PII compost on one side and 11.5 g PIII compost on the other side from which the PII compost was colonized. Growth in boxes was done using 1 kg (30 × 20 × 22 cm boxes), 8 kg (40 × 60 × 20 cm boxes), or 16 kg (40 × 60 × 22 cm boxes) PII-end compost that was inoculated with 8, 64, or 75 gr of spawn, respectively. The 1 and 8 kg compost boxes that were used to monitor temperature dynamics during PIII were covered with plastic film containing 50 3-mm-wide holes m^-2^. The 16 kg compost boxes were used to monitor temperature dynamics both during PIII and PIV under cultivation conditions described previously (Vos, Jurak, *et al*., 2017).

### Quantification of CO_*2*_, *O*_*2*_, *temperature and growth*

Correlation of CO_2_ production and O_2_ consumption was done using a respirometer (Biometric Systems, Germany) with optical CO_2_ and O_2_ sensors. In short, CO_2_ and O_2_ was measured using biological triplicates at t=0 and t=2.5 h, after which the air in the flask was refreshed. This was repeated 170 times during a 425 h period. Data were converted to molar amounts of produced CO_2_ and consumed O_2_, correcting for temperature and pressure.

Growth in compost was measured by placing 12-cm-square Petri dishes in a flatbed scanner (Epson V300 / V370, Seiko Epson Corporation, Shinjuku, Tokyo, Japan). Compost was scanned every hour. These images were produced in 16-bit grayscale intensity at 2400 dpi resolution (10.6 µm pixel size). The grey values were quantified in a 3.8 × 0.5 cm window around a Pt100 (class B) temperature sensor probe (1.83 × 6 mm; Sensing Devices LLC, Lancaster, USA) that had been placed inside the 2 cm layer PII compost in the middle of the square Petri dish before placing the dish in the scanner. Temperature was logged by connecting the temperature probe to a real-time clock (RTC) DS1307 module, a micro-SD module, a MAX31865 RTD module, and an Arduino Nano according to manufacturer’s instructions. Temperature was measured every 5 minutes, while the mean grayscale intensity within the temperature probe area was quantified over time using ImageJ. The scanner was covered with a 5 cm thick walled polystyrene box to minimize the effect of the incubator hysteresis.

Arduinos (https://www.arduino.cc/) equipped with temperature (DS18B20; Dallas Semiconductor, USA) or CO_2_ (MG811; Sandbox Electronics, Finland) sensors were used to measure and correlate changes in temperature and CO_2_ levels in box experiments. To this end, the temperature sensor was placed 2 cm below the compost surface, while the CO_2_ sensor was placed 5 cm above the compost.

Vegetative incompatible strains (A15/Bisp015) and vegetative compatible strains (A15/A15 and Bisp015/Bisp015) were grown together in duplo in a 40×60 cm compost box filled with 16 kg compost. The compost was divided in 4 areas, 2 of which were inoculated with A15 and two with Bisp015 in diagonal positions. Two thermocouple probes (Type T; Copper/Constantan) were placed per separate area (8 probes in total). Temperature changes were recorded using a DT85 Series 2 Data logger (Cambeep, Cambridge, United Kingdom), every 10 min starting on day 1 of cultivation and ending at day 18.

### RNA isolation

RNA was isolated from 7.5 gr colonized compost of box cultures (30 × 20 × 22 cm) after 130-145 h of incubation (Figure S5). Samples (biological triplicates) were taken from the immediate surroundings of a temperature sensor during and after a burst and in between bursts (i.e., 0.5, 4.5, and 8.5 h after a burst started, respectively). Compost was crushed using mortar and pestle under liquid nitrogen and ground to a fine powder for 1 min at 30 Hz using a tissuelyser II with a steel grinding jar pre-cooled with liquid nitrogen (Qiagen, Venlo, Netherlands). RNA was extracted using a modified method described by Patyshakuliyeva et al. (2014). For each sample, 200 mg compost powder was transferred to 7 ml extraction buffer (4 M guanidinium thiocyanate, 25 mM sodium citrate, pH 5.0, 0.5 % N-lauroyl sarcosine, 0.1 M β-mercaptoethanol), shaken vigorously, and incubated for 10 min at RT. Next, 7 ml of chloroform:isoamyl alcohol (24:1) was added, mixed by vortexing, and incubated for 2 min. Phases were separated by centrifugation for 15 min at 4500 g and 6.5 ml of the aqueous phase was transferred to an ultracentrifugation tube (Ultra-Clear, open-top; Beckman coulter) filled with a 2 ml cushion of 5.7 M CsCl in 25 mM sodium citrate, pH 5.0. The tube was topped with 7 ml 0.8 M guanidine thiocyanate, 0.4 M ammonium thiocyanate, 0.1 M sodium acetate, pH 5 (TS buffer). RNA was pelleted by centrifugation for 24 h at 4 °C and 104.000 g in a SW28.1 swing out rotor (Beckman Coulter, Brea, CA, USA). RNA was taken up in 100 µl H_2_O and 500 µl TS buffer was added. RNA was precipitated with 0.8 volume isopropanol, washed with 70 % ethanol, and further purified using a Qiagen RNeasy Mini Kit. RNA quality and quantity was assessed using gel electrophoresis and a Qubit Fluorometer.

### RNA sequencing and analysis

Strand-specific mRNA-seq libraries for the Illumina platform were generated and sequenced at BaseClear BV (Leiden, The Netherlands). High-quality total RNA (checked and quantified using a Bioanalyzer (Agilent, Santa Clara, CA, USA) was used as input for a dUTP library preparation method (Parkhomchuk *et al*., 2009; Levin *et al*., 2010). For dUTP library preparation the mRNA fraction was purified from total RNA by polyA capture, fragmented and subjected to first-strand cDNA synthesis with random hexamers in the presence of actinomycin D. The second-strand synthesis was performed incorporating dUTP instead of dTTP. Barcoded DNA adapters were ligated to both ends of the double-stranded cDNA and subjected to PCR amplification. The resulting library was quality checked and quantified using a Bioanalyzer (Agilent). The libraries were multiplexed, clustered, and sequenced on an Illumina HiSeq 2500 (HiSeq Rapid SBS v2 chemistry) with a single-read 50 cycles sequencing protocol and indexing. At least 10 million 50 bp single reads were generated per sample.

FASTQ sequence files were generated using bcl2fastq2 version 2.18. Initial quality assessment was based on data passing the Illumina Chastity filtering. Subsequently, reads containing PhiX control signal were removed using an in-house filtering protocol. In addition, reads containing (partial) adapters were clipped up to minimum read length of 50 bp. Remaining reads were subjected to a second quality assessment using the FASTQC quality control tool version 0.11.5 (http://www.bioinformatics.babraham.ac.uk/projects/fastqc/) and deposited in NCBI GEO with accession number GSE124976. HISAT version 2.1.0 (Kim *et al*., 2015) was used to align sequence reads to the Agabi_varbisH97_2 version of the *A. bisporus* H97 genome (Morin *et al*., 2012), which was obtained from MycoCosm (Grigoriev *et al*., 2014). Cuffdiff (version 2.2.1), which is part of Cufflinks (Trapnell *et al*., 2010), was used to identify reads mapping to predicted genes and to identify differentially expressed genes. The bias correction method was used while running Cuffdiff (Roberts *et al*., 2011). Cuffdiff normalizes the expression level of each predicted gene to fragments per kilobase of exon model per million fragments (FPKM). In addition to Cuffdiff’s requirements for differential expression the following requirements were applied: a ≥2-fold change and a minimal expression level of 10 FPKM in at least one of the samples. Quality of these results was analyzed using CummeRbund (Goff *et al*., 2013). All expression data can be found in Dataset S4.

### Functional annotation

Conserved protein domains were predicted using PFAM version 31 (Finn *et al*., 2016) and mapped to the corresponding gene ontology (GO) terms (Hunter *et al*., 2009). Proteases were predicted using the MEROPS database (Rawlings *et al*., 2016) with a blastp E-value cut-off of 10^−5^. Secretion signals and transmembrane domains were predicted using SignalP 4.1 (Petersen *et al*., 2011) and TMHMM 2.0c (Krogh *et al*., 2001), respectively.

The previously identified *A. bisporus* CAZYs and proteins involved in carbon metabolism (Patyshakuliyeva *et al*., 2013, 2015) were further annotated and supplemented as described in Supplementary Text 1. Over- and under-representation of functional annotation terms in sets of differentially regulated genes were identified using the Fisher Exact test. The Benjamini-Hochberg correction was used to correct for multiple testing using a p-value < 0.05.

## Results

### Respiratory bursts in compost

The CO_2_ production and O_2_ consumption of PII-end compost inoculated with *A. bisporus* variety Sylvan A15 (A15) was monitored using a respirometer. Non-inoculated and sterilized PII-end compost served as a control. The CO_2_ production and O_2_ consumption were determined every 2.5 h during a 425 h measurement period that started after a 5-days pre-incubation at 25 °C. Total CO_2_ produced per gram of sterile compost, non-inoculated compost, and inoculated compost was 52, 227, and 831 µmol during the 425 h period, respectively (Figure 1A). The CO_2_ production rate decreased over time from 1 to 0.3 µmol h^-1^ g^-1^ in compost without *A. bisporus* A15, while ≤ 0.23 µmol CO_2_ h^-1^ g^-1^ compost was released from sterile compost (Figure 1B). In contrast, CO_2_ production in compost with *A. bisporus* A15 increased from 1.2 to 2.6 µmol h^-1^ g^-1^ substrate during the first 180 h and then decreased to 1.3 µmol h^-1^ g^-1^ compost. This was accompanied by distinct respiratory bursts of 2.5-5 h (Figure 1B). The CO_2_ production rate during the bursts increased up to 3.5 fold as compared to the intermediate periods. These intermediate periods had an initial length of 13 h and increased to 20 h. The interval became irregular as illustrated by an increase to 50-90 h after 200 and 240 h, respectively. No respiratory bursts were observed in the absence of *A. bisporus* A15 but they were produced in sterilized compost inoculated with this fungus (Figure S1). However, growth of *A. bisporus* A15 in sterilized compost was slow coinciding with low CO_2_ production.

**Figure 1.**
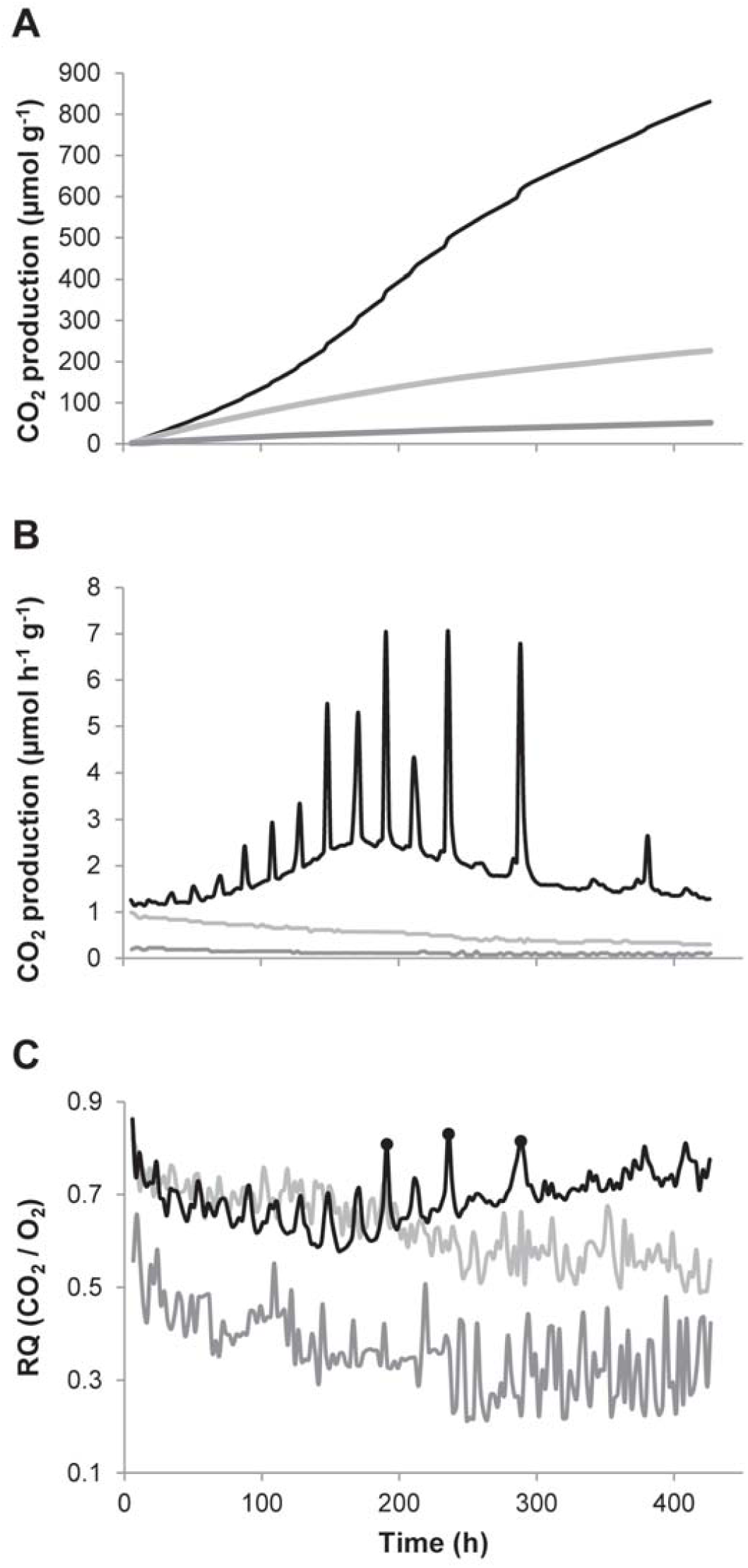
Cumulative CO_2_ production (A), CO_2_ production rate (B), and respiratory quotient (C) by sterile compost (dark grey), compost without *A. bisporus* (light grey), and compost inoculated with *A. bisporus* (black). O_2_ levels were below 17.97 % (the limit at which O_2_ could be measured) at time points indicated with a black dot (C), resulting in an underestimation of the actual O_2_ consumption and therefore overestimation of the RQ.

The respiratory quotient (RQ; mol CO_2_ released / mol O_2_ consumed) in non-inoculated compost decreased from 0.8 to 0.55 during the 450 h measurement period (Figure 1C), while it decreased in sterilized compost from 0.6 to 0.3. During the colonization of *A. bisporus* A15 the maximal RQ decreased from 0.8 to 0.65 during the first 150 h period and then increased to 0.8 again in the following 350 h (Figure 1C). At three time points the O_2_ levels were below 17.9 % resulting in an underestimation of the actual O_2_ consumption and overestimation of the RQ (Figure 1C, black dots). The RQ was up to 0.13 higher during bursts as compared to the pre-burst minimum of 0.58.

CO_2_ production correlated with an increase in compost temperature. Temperature increase was detected 30 min after the start of a respiratory burst, peaked 45-90 min later, and took 2-8 h to normalize to pre-peak temperature (Figure S2). Hence, monitoring compost temperature can be used to monitor respiratory bursts. This was adopted to monitor respiratory bursts during PIII and PIV of a semi-commercial *A. bisporus* A15 cultivation. Respiratory bursts were observed between day 10 and 16 of PIII (Figure S3) and followed a similar pattern as observed in small scale lab cultivations. They were also observed during colonization of casing and the 1^st^ flush but not during the 2^nd^ flush.

### Synchronization of respiratory bursts is mediated by hyphal fusion

The respiratory bursts observed during colonization of compost by *A. bisporus* A15 are suggestive of synchronized mycelial action in the compost. This synchronization was further characterized using 6 temperature sensors that had been placed 15-25 cm apart in a 40 × 60 cm box containing 8 kg inoculated PII-end compost. *A. bisporus* A15 was allowed to colonize the compost for 8 days at 25 °C before temperature measurements were started. Respiratory bursts occurred during the first 400 h of a 700 h measurement period (Figure 2A). They resulted in a temperature increase of up to 2 °C in a 1.5-3 h period. Initially, the time delay between all sensors measuring a burst was 170 min (Figure 2B). This decreased to 50 min between 33 and 88 h after the measurements had been started (Figure 2B), showing that heat production by the mycelium synchronized over at least 50 cm (the largest distance between the sensors). No pattern was observed in the order sensors measured a burst. After 120 h the time delay of the sensors picking up a burst increased to 116 min, while after 198 h not all sensors picked up a burst anymore with one exception at 218 h. After this time point bursts became erratic.

**Figure 2.**
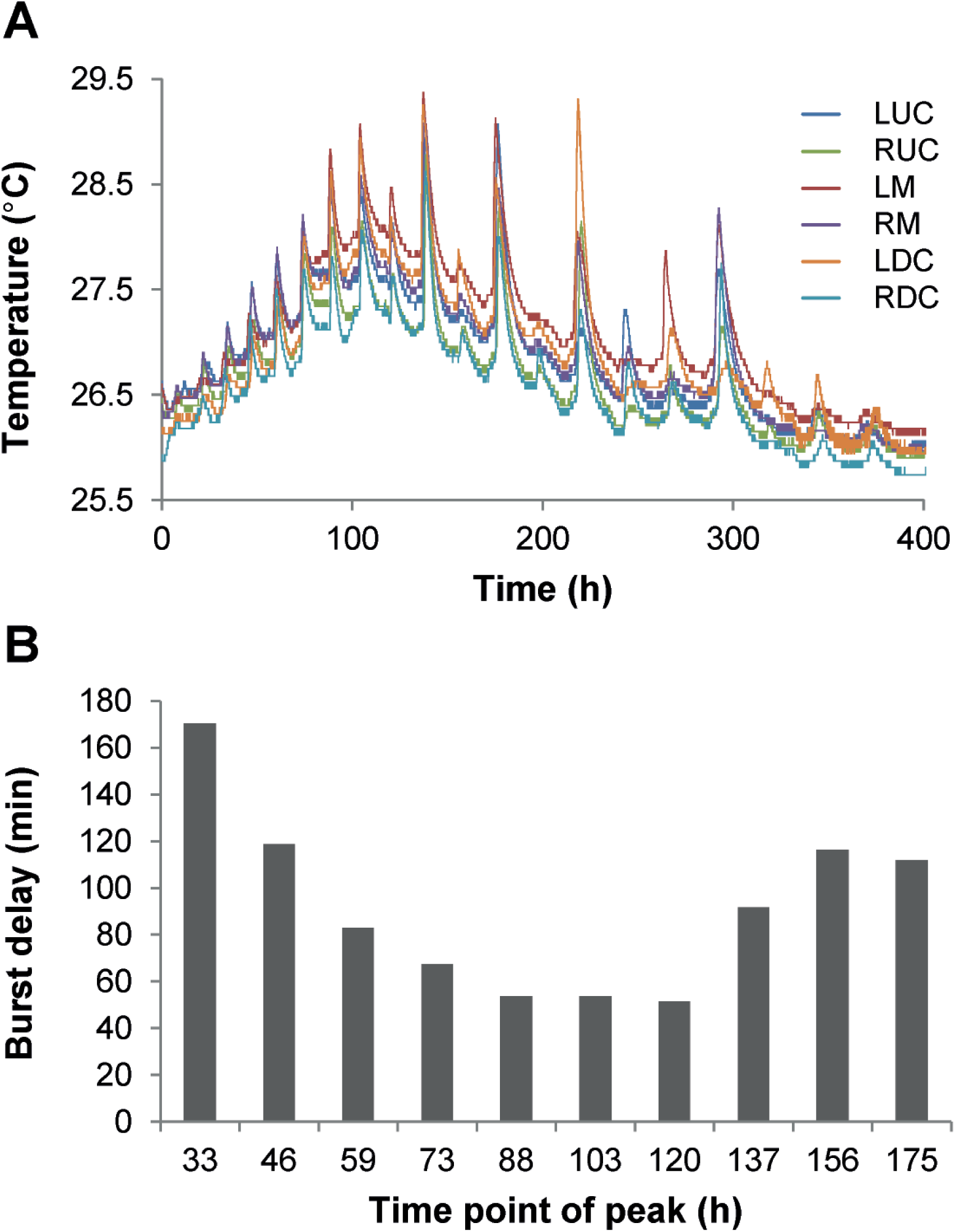
Compost temperature measured by 6 sensors spaced 15-25 cm apart in a 40 × 60 cm box (A) and the maximum peak delay between the sensors during a burst (B). LUC = left upper corner, RUC = right upper corner, LM = left of the middle, RM = right of the middle, LDC = left bottom corner, RDC = right bottom corner.

To obtain information on the nature of the synchronization in compost two aliquots of 1 kg *A. bisporus* A15 inoculated PII-end compost were placed next to each other (i.e. physically touching each other), 3 cm apart, or were separated by aluminium foil. In the latter case, transfer of heat could occur but hyphae of the two parts could not interact. The temperature in the centre of both parts of compost was monitored for 300 h after 8 days of culturing (Figure 3). The average time between sensors in the two parts of compost picking up a burst was lower when the parts were in physical contact (38 min) when compared to those placed 3 cm apart or separated by aluminium foil (3 h 50 min and 4 h 33 min, respectively; Mann-Whitney U, p < 0.05). This suggests that interaction of mycelium is required for the observed synchronization.

**Figure 3.**
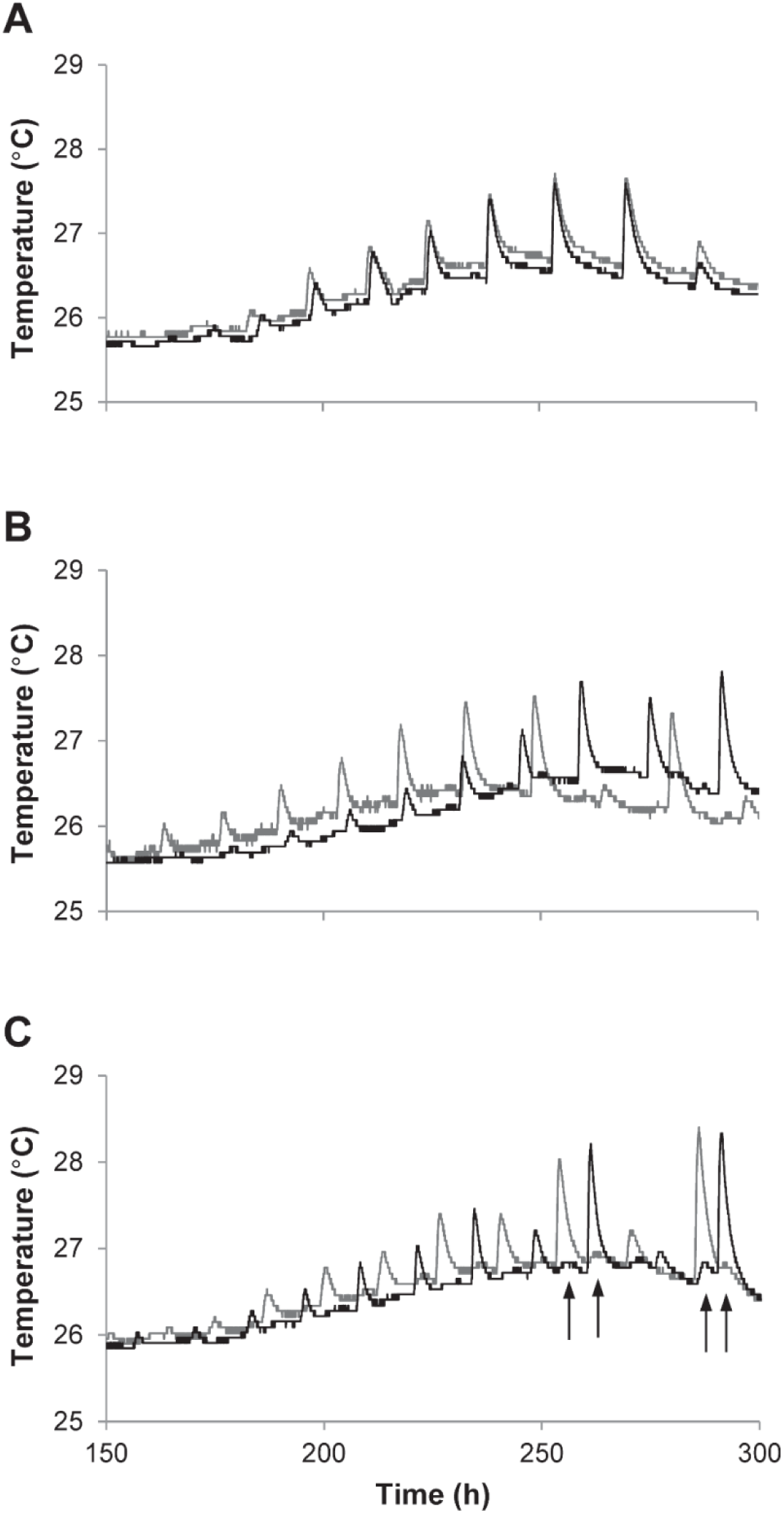
Typical temperature profiles of two parts of compost (dark and light grey) pressed together (A), placed 3 cm apart (B), or separated by aluminium foil (C). Arrows indicate where heat transfer through the aluminium foil was detected by the temperature sensor. Each experiment was performed 5 times. The average difference between two temperature sensors picking up a burst (i.e. the time delay of a burst) for each condition was tested using Mann-Whitney U, p < 0.05.

Hyphae of *A. bisporus* colonies that are vegetatively compatible can fuse to form a larger colony (O’Connor *et al*., 2020). The use of two incompatible strains allows to distinguish between proximity of hyphae or hyphal fusion being required for synchronization of bursts over large distances. To this end, compost was divided in 4 areas of which two were inoculated with *A. bisporus* A15 and two with *A. bisporus* Bisp015 in diagonal positions in a 40 × 60 cm box. The mycelium of *A. bisporus* A15 anastomoses with hyphae of other *A. bisporus* A15 colonies forming a single mycelium (Figure S4). However, *A. bisporus* A15 does not anastomose with hyphae of *A. bisporus* Bisp015 as these strains show a vegetative incompatibility, i.e. do not fuse likely due to different alleles in vegetative incompatibility genes (O’Connor *et al*., 2020) (Figure S4). Temperature sensors placed in an area containing only 1 of both strains synchronized, while no synchronization of bursts was observed between areas containing A15 and areas containing Bisp015 (Figure 4A and 4B). This shows that anastomosis is required to synchronize the activity of the mycelial colony.

**Figure 4.**
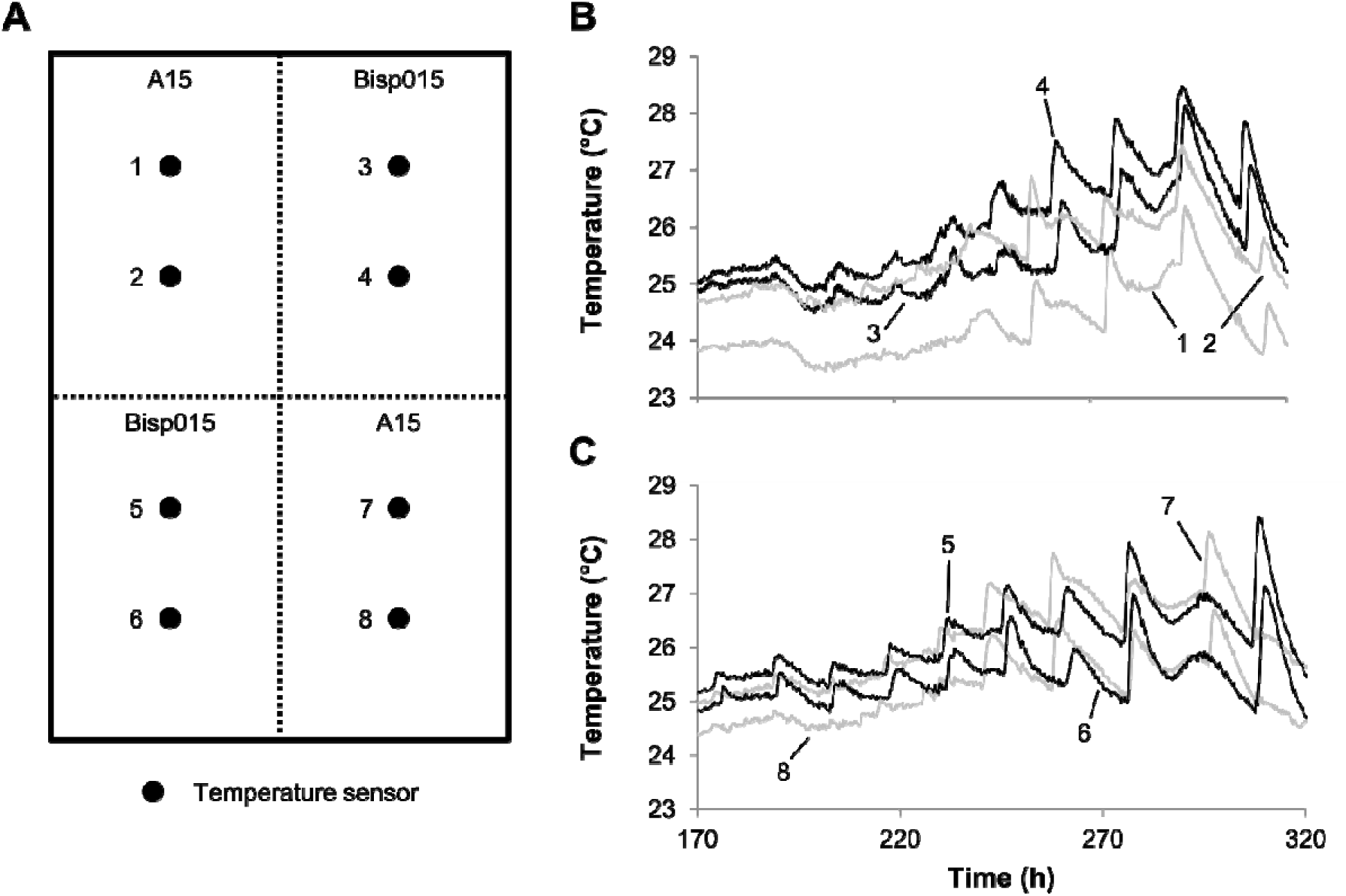
Experimental setup (A) and typical temperature profile of compost divided in 4 areas, 2 of which were inoculated with A15 (B and C, black lines) and two with Bisp015 (B and C, grey lines) in diagonal positions in a box. Numbers in panel B and C relate to the location of temperature sensors in panel A. The temperature profiles of A15 and Bisp015 in panel B and C are representative of 2 independent experiments.

### Respiratory bursts occur in sub-peripheral zones of the mycelium

In all previous experiments the inoculum of *A. bisporus* was homogeneously distributed throughout the compost. Therefore, it is not possible to distinguish if bursts occur at the hyphal front of a colony or if older parts of the mycelium contribute to these bursts. To distinguish between these options, growth of *A. bisporus* A15 containing PIII compost into fresh PII compost in a square Petridis was monitored using a flatbed scanner and a temperature sensor. Biomass was distinguished by increased whitening of the compost and the temperature sensor was placed in the middle of the PII compost to monitor the bursts in a moving hyphal front. The linear growth rate of the hyphal front was 8.8 mm d^-1^. After 163 h a small temperature increase was observed when the mycelial front had approached the temperature sensor to 8.1 mm (Figure 5). This was followed by bursts starting 9 h after the mycelium had reached the temperature sensor; the hyphal front had grown 3.6 mm past the sensor at this moment. Bursts were separated by 23-36 h periods. Notably, bursts continued after the mycelial front had passed the probe up to 5.7 cm. At this moment, hardly any biomass was produced anymore at this location as shown by the low increasing of whitening of this area. Moreover, respiratory burst were not accompanied by increased growth as the increase and decrease in grayscale value around bursts was caused by condensation of water and its evaporation. Together, the data indicate that the whole colony of *A. bisporus* (i.e. the hyphal front and older parts of the colony) contribute to the respiratory bursts observed in previous experiments.

**Figure 5.**
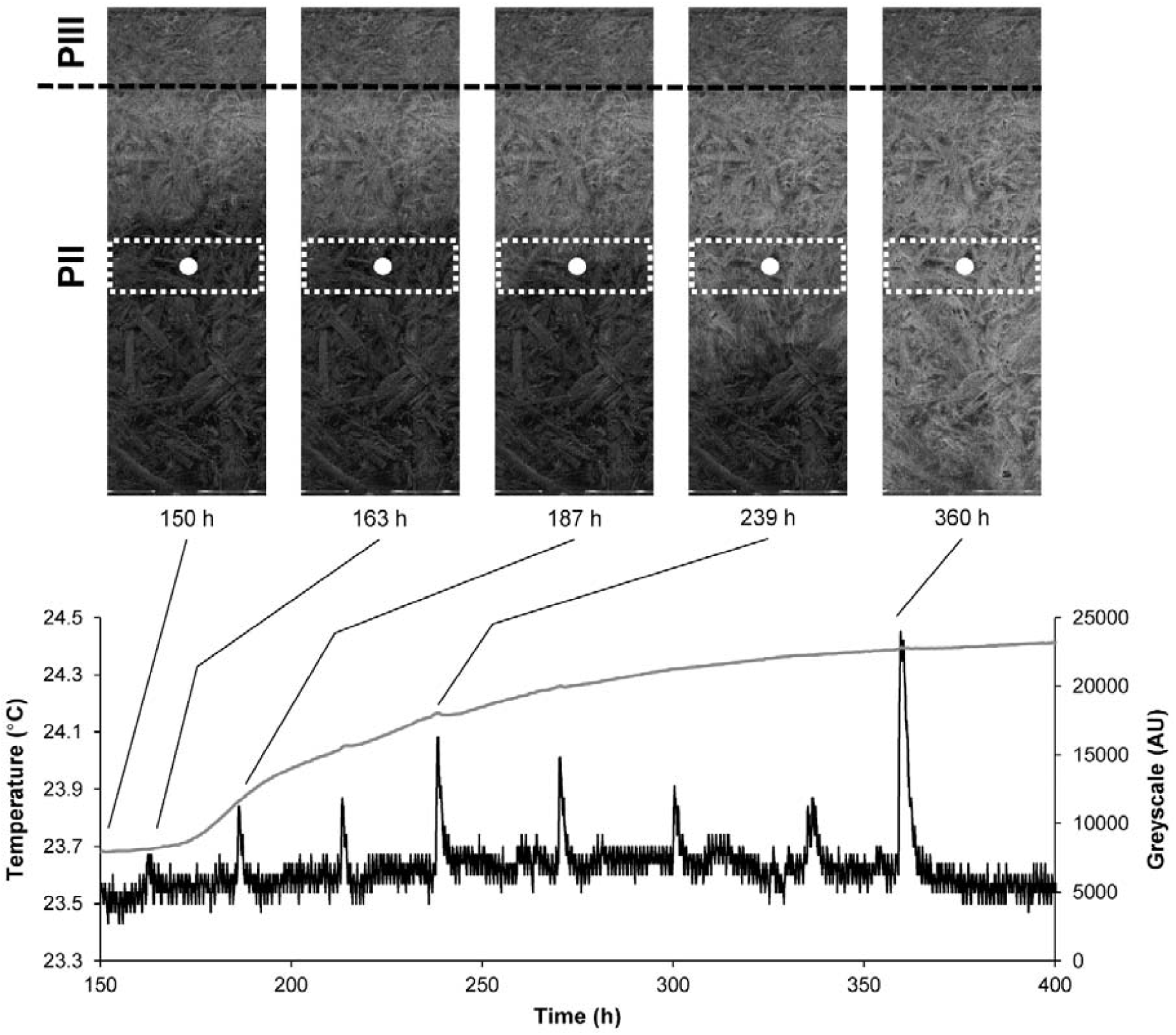
Compost temperature (black line) and mycelial growth expressed as grey value (grey line, based on the area in the white dashed boxes) during growth of *A. bisporus* from PIII compost into fresh PII compost (border indicated by the dashed line). Scans of the compost after 150, 163, 187, 239, and 360 h are shown with the position of the temperature sensor indicated (white dot). These data are representative of 2 independent experiments.

### Reorganization of gene expression during respiratory bursts

The physiological characterization of vegetative growth of *A. bisporus* revealed a burst-like phenomenon not previously observed in fungi growing in complex substrates. RNA was isolated from biological triplicates during, after, and in between bursts (0.5 h, 4.5 h, and 8.5 h after a burst started, respectively) to investigate the process underlying these bursts (Figure 6A). Out of the 10438 genes encoded by the *A. bisporus* genome, 4731 were differentially expressed (i.e. ≥ 2-fold up or down-regulated, with an fpkm > 10 in at least 1 of the 3 conditions and a q-value < 0.05; Figure 6B). A total of 197 GO terms and 69 Pfam domains were over- and / or under-represented in sets of up- or down-regulated genes (Dataset S1). Cell cycle (i.e. cell division and / or nuclear division) and translation related GO terms were overrepresented and underrepresented, respectively, in upregulated genes after a burst relative to during a burst, while GO terms related to transcription and the cell membrane were overrepresented in genes upregulated after a burst when compared to the other two samples. The GO term related to metabolic processes was overrepresented in genes upregulated during the bursts relative to the inter-burst samples and in downregulated genes after a burst relative to the other two samples. Furthermore, cell wall related GO terms were overrepresented in genes downregulated during a burst when compared to the inter-burst samples. Proteins with a predicted secretion signal were overrepresented in both upregulated and downregulated genes during a burst compared to the inter-burst samples. In addition, these proteins were overrepresented in downregulated genes after a burst relative to the other two samples. The differential expression of cyclins after a burst points to a synchronized cell cycle (Supplementary Text 2). It is not clear which functions these cyclins have; i.e. whether they are involved in nuclear division or in growth.

**Figure 6.**
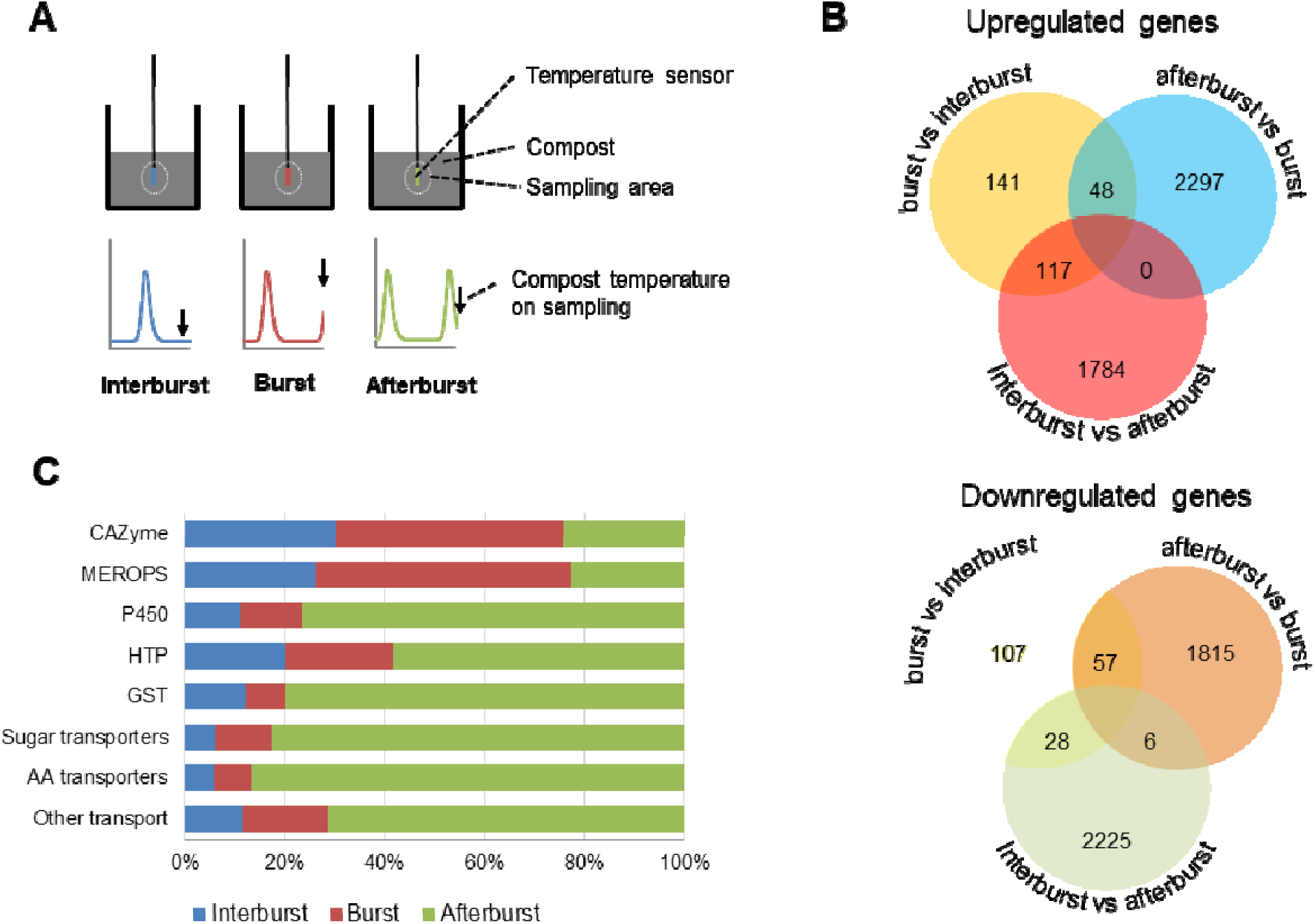
Schematic representation of experimental setup for the transcriptome analysis of respiratory bursts (A) and analysis of differentially expressed genes. Samples were harvested in between bursts (end of blue line), during bursts (end of red line), or after a burst (end of green line). For each point, RNA was isolated from 3 biological replicates. Up-(B, top panel) and down-(B, lower panel) regulated genes before a burst, during a burst, and after a burst (i.e. ≥ 2-fold up or down-regulated, with an fpkm > 10 in at least 1 of the 3 conditions and a q-value < 0.05). Relative abundance of FPKM of differentially expressed genes annotated as CAZYs, MEROPS proteases, P450 enzymes, heme-thiolate peroxidases (HTP), gluthatione-S-transferase (GST), sugar transporters, amino acid transporters, and other transporters (C).

Expression of CAZYs and proteases was increased during a burst, while enzymes with a potential role in metabolism of lignin derived products (P450, Heme-thiolate peroxygenase [HTP], β-etherase), and sugar, amino acid, and other transporters were highly expressed after a burst (Figure 6C). Putative sugar transporter genes (Pfam 00083) were overrepresented in the downregulated genes during the inter-burst relative to after a burst (Dataset S1). Indeed, the majority of differentially expressed sugar transporters were upregulated after a burst (Figure 6C; Dataset S7; 19 out of 33). In addition, differentially expressed amino acid / peptide transporter genes were mostly upregulated after a burst (20 out 26; Figure 6C; Dataset S7; Pfam 03169, 00854, 01490, 00324, and 13520). From the remaining 50 differentially expressed transporter genes (annotated with Pfam 07690; Figure 6C; Dataset S7) 36 were upregulated after a burst. These genes included putative nicotinic acid and thiamine transporter genes and transporter genes putatively involved in secretion of toxins (Pfam 06609; Alexander *et al*., 1999). The combination of transporters and secreted proteins being differentially expressed points to changes in the extracellular availability of free sugars. Therefore, the CAZYs were analysed in more detail. Expression values of each differentially expressed CAZY family were summed to identify general patterns in their expression (Figure 7).

**Figure 7.**
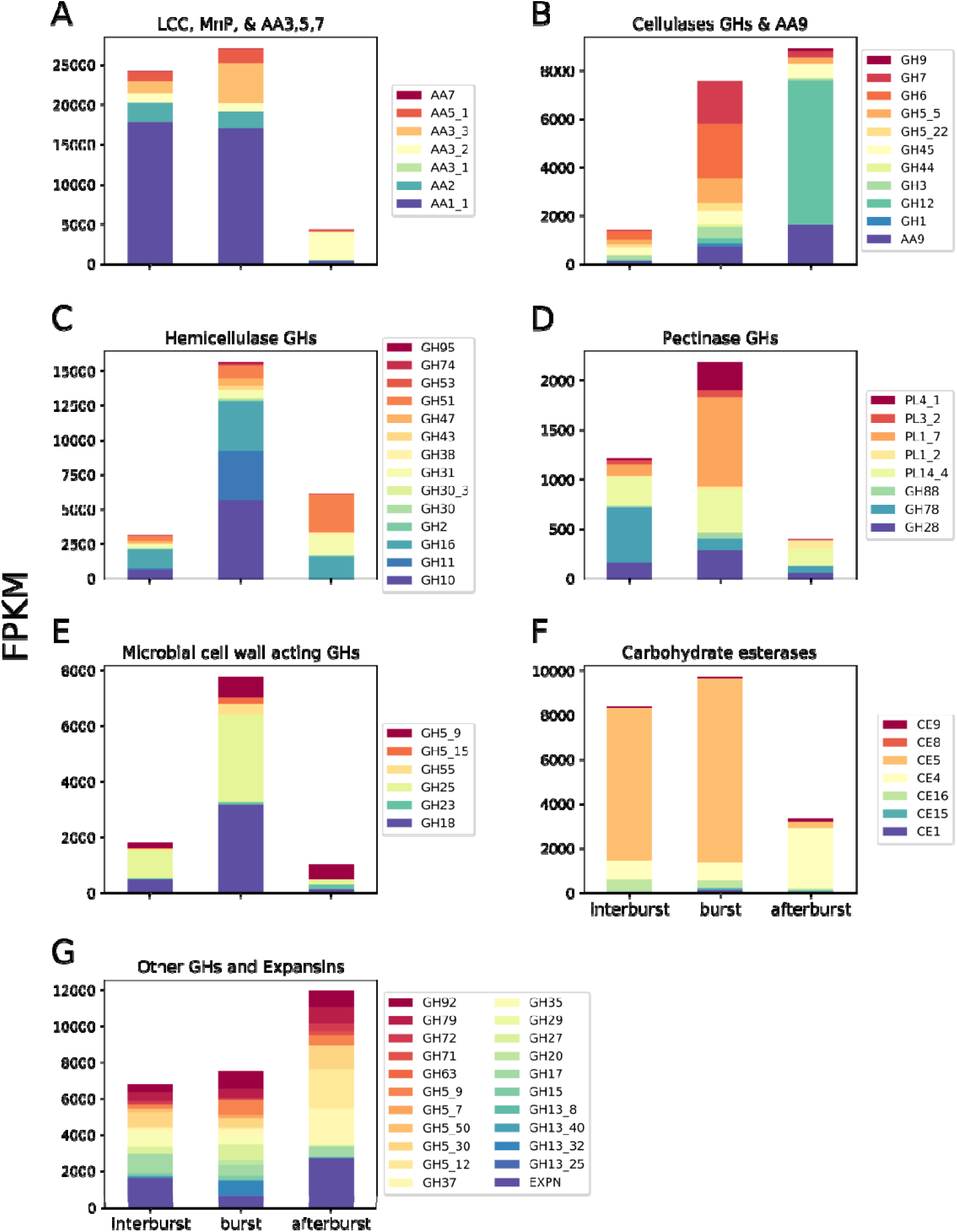
Expression values (FPKM) of differentially expressed classes of CAZYs related to lignin degradation (A), cellulose degradation (B), hemicellulose degradation (C), pectin degradation (D), microbial cell wall degradation and remodeling (E), carbohydrate esterases (F), other GHs and expansins (G). Bars represent expression values before (inter-burst) during (burst) and after bursts (after burst).

### Downregulation of the ligninolytic system after bursts

MnPs and LCCs (AA2 and AA1_1) were highly expressed before and during bursts but downregulated up to 367 fold for the most highly expressed LCCs after bursts (Figure 7A; Dataset S2 and S3). Similarly, glyoxal oxidases (GLOX; AA5_1) and alcohol oxidases (AO; AA3_3) were down-regulated (3.5-194 fold) after a burst and upregulated 2.3-58 fold in between bursts. Putative aryl alcohol oxidases, glucose 1-oxidases, and pyranose dehydrogenases (AA3_2) were upregulated after a burst and downregulated in the interburst samples. Expression of fatty acid desaturases (Protein ID 194591 and 193825; Dataset S4) peaked after a burst, being upregulated 13.6-243 fold compared to a burst and the inter-burst. These data show that genes encoding the ligninolytic machinery are actively expressed before and during bursts, while they are downregulated after bursts with the exception of fatty acid desaturases.

### Polysaccharide deconstruction CAZYs are upregulated during respiratory bursts

Cellulase genes (Figure 7B; Dataset S2 and S3) were upregulated during and after bursts relative to in between bursts. Putative endo- and exo-acting cellulases like GH5_5, GH6, GH7, GH45, and the LPMO AA9 dominated the expression of cellulases during bursts while the endoglucanase GH12 genes together with AA9 and GH45 were dominantly expressed after bursts. Hemicellulases peaked in expression during bursts and were dominated by putative endo-acting hemicellulases like GH10, GH11, and GH16 while after bursts GH16, GH31, and GH51 were dominantly expressed (Figure 7C). Similar to the ligninolytic system pectinases were downregulated after bursts (Figure 7D). In between bursts the GH78 and the PL14_4 pectinase families were dominantly expressed while the PL4_1, PL1_7, PL14_4, and GH28 pectinase families were most highly expressed during bursts. Glycoside hydrolase families acting on microbial cell walls were upregulated during bursts (Figure 7E) and their expression was dominated by lysozyme and chitinase families GH18 and GH25, and putative β-1,3-glucanases and β-1,6-glucanases (GH5_9, GH5_15, and GH55). Putative cutinases of the carbohydrate esterase family CE5 were dominantly expressed in between and during bursts (Figure 7F; Dataset S2 and S3). The CE5 family was downregulated after bursts while the putative chitin deacetylases of the CE4 family were upregulated after a burst. Finally, expression of expansins was lower during bursts compared to the samples in between and after bursts (Figure 7G). From these data it is clear that genes involved in degradation of hemicellulose and microbial cell walls are primarily expressed during bursts while cellulases are also expresses after a burst.

### Sets of protease genes are upregulated during and after respiratory bursts and in inter-burst periods

Upregulation of amino acid and peptide transporters after bursts point to increased extracellular availability of theses nutrients at this stage. MEROPS protease genes were enriched in the upregulated genes during a burst, in the downregulated genes after a burst and the upregulated genes during the inter-burst (Dataset S1). Specifically, sets of protease genes were expressed during, after and in between bursts (Dataset S5). For example, 19 out of 23 and 49 out of 54 protease genes upregulated during and after a burst, respectively, were most highly expressed at that sample point. The highly expressed serine protease gene SRP1 (Burton *et al*., 1997; Heneghan *et al*., 2016) (Protein ID 194648) that is implicated in nitrogen acquisition from humic-rich substrates was upregulated 3.5 fold during a burst and 372 fold downregulated after this event. Together, this suggests a highly differentiated role for proteases in vegetative growth of *A. bisporus*.

### Expression of metabolic genes support a repeated reorganization of fungal metabolism

A total of 15 out of 59 genes involved in glycolysis, the pentose phosphate pathway (including genes involved in conversion of pentoses to D-xylulose), and the galactose pathway changed expression more than 5 fold during cycling (Dataset S8). For example, the pyruvate kinase (EC 2.7.1.40), ribose-phosphate pyrophosphokinase (EC 2.7.6.1) and ribulose-phosphate 3-epimerase (EC 5.1.3.1) genes were upregulated 37, 10, and 16 fold after a burst, respectively. Furthermore, 7 out of 21 genes involved in the TCA cycle changed expression ≥ 5 fold of which 4 were downregulated after a burst and 2 were upregulated after a burst. A predicted trehalose phosphorylase (EC 2.4.1.231/2.4.1.64) and mannitol-1-phosphate dehydrogenase (EC 1.1.1.17) gene were upregulated 3.8 and 13 fold, respectively, after a burst. This indicates that the metabolism of *A. bisporus* is regulated in a cyclic way during most of the vegetative growth in compost.

The secreted oxalate decarboxylase (Kathiara *et al*., 2000) (EC 4.1.1.2) and intracellular formate dehydrogenase (EC 1.17.1.9) genes were downregulated 5.3 and 105 fold, respectively, after a burst and stably expressed in between and during bursts. Two glyoxalase genes (Pfam 00903) and a putative glyoxylate reductase gene (EC 1.1.1.26) were also downregulated after a burst (5.6-39 fold), while the putative glyoxylate reductase gene and a glyoxalase gene were upregulated before and during a burst (2.3-17.9 and 2.2-2.5 fold, respectively). This shows that genes involved in oxalate metabolism are downregulated after bursts.

## Discussion

Growth of filamentous fungi in complex substrates is associated with respiration (i.e. production of CO_2_ and O_2_ consumption) and heat production, yet these variables are rarely measured. Here, a respirometer and temperature electrodes were used to study the vegetative growth of *A. bisporus* in compost revealing bursts of temperature and respiratory activity. Respiratory cycles with periods varying between 7.5 and 22.5 h (i.e. distinct from a circadian type of rhythm) have been observed during colonization of malt extract and wood blocks by the brown rot fungi *Neolentinus suffrutescens* (*Lentinus lepideus*), *Gloeophyllum trabeum (Lenzites trabea), Postia placenta (Poria monticola), and Coniophora puteana* (Damaschke and Becker, 1966; Smith, 1973). This indicates that respiratory patterns are widespread in basidiomycetes. However, the increase in respiration during bursts produced by *A. bisporus* is more dramatic (up to 3.5-fold) compared to the wave-like respiratory patterns of brown rot fungi (up to 0.3-fold). Bursts of *A. bisporus* became increasingly synchronized over a distance of at least 50 cm and required anastomosis of hyphae as the burst produced by incompatible strains A15 and Bisp015 did not synchronize. Older parts of the mycelium participated in bursts as they were observed until the outer region of the mycelium had passed the temperature sensor 5.7 cm. Thereafter, the mycelium had reached the edge of its substrate and bursts were no longer observed. Growth of the hyphal front may, therefore, be important for the occurrence of bursts in older parts of the mycelium.

Compost temperature increased up to 2 °C in 1.5 to 2 h during bursts corresponding to the release of up to 5 kJ energy kg^-1^ compost when assuming a water content of 60 %. Thornton’s Rule shows that O_2_ consumption can fully explain the temperature increase observed in compost (Thornton, 1917; Hansen *et al*., 2004) (Supplementary text 3; Table S1, S2). The decrease in RQ from 0.8 to 0.65 in the period where respiratory bursts are most regular (Figure 1C, first 150 h) fits with the formation of oxidized compounds like oxalic acid that are formed around hyphae of *A. bisporus* during its vegetative growth (Atkey and Wood, 1983) and preferential lignin mineralization during PIII (Kabel *et al*., 2017). Lignin mineralization accounts for approximately 84 % of the total heat production during PIII (Supplementary text 3; Table S3, S4), while the total O_2_ consumption during bursts is 21 % of total O_2_ consumption (Table S1). Therefore, lignin degradation cannot occur solely in bursts as oxygen consumption does not match what is expected from lignin loss.

Transcriptomic data, obtained in between, during, and after bursts, supports the view that *A. bisporus* cycles through degradation of lignin, (hemi-) cellulose and microbial biomass, and uptake of breakdown products using a synchronized vegetative growing mycelium. The ligninolytic system of *A. bisporus* is active before and during these bursts while (hemi-) cellulolytic genes and genes encoding microbial biomass degrading activities are upregulated during and after this event. The active expression of sugar and amino acid transporters after a burst indicates that carbon influx occurs at this time. Indeed, genes involved in glycolysis and the pentose phosphate pathway were upregulated (up to 37 fold) after a burst. In addition, expression of trehalose phosphorylase and mannitol-1-phosphate dehydrogenase genes was increased after bursts too (up to 13 fold). Therefore, trehalose and mannitol may not only play a role in mushroom formation (Hammond and Nichols, 1976) but also in carbon storage and transport within the mycelial network.

The increase in respiration and the oscillations of the RQ around the time bursts occur (e.g. from 0.58 to 0.71) show that there is increased mineralization of more oxidized substrates during a burst. This could be explained by the consumption of sugars (RQ = 1) but also mineralization of oxalic acid (RQ = 4). This link between bursts and oxalate metabolism is supported by downregulation of putatively secreted oxalate decarboxylase and formate dehydrogenase after bursts (up to 105 fold). Thus, the bursts and RQ oscillations could relate to mineralization of extracellular oxalic acid. Free energy conservation in oxalate (or sugar) metabolism produces ATP that would be required for synthesis of secreted proteins like CAZYs during bursts. Protein production is an energy intensive process (Stouthamer, 1973). Hence, protein synthesis could partly explain the temperature increase (O_2_ consumption) during bursts. Protein secretion could be facilitated by formation of new (small) branching hyphae or cell wall remodeling of older mycelium (Wosten *et al*., 1991; Krijgsheld *et al*., 2013). Furthermore, a burst was observed in the 1^st^ flush of a semi-commercial cultivation but not in the 2^nd^ flush. Possibly relating to previous observations where low expression of CAZYs was found in samples taken during the 2^nd^ flush (Patyshakuliyeva *et al*., 2015).

In line with downregulation of the ligninolytic system, MnP, LCC, glyoxal oxidases, and alcohol oxidase genes were downregulated up to 194-fold after a burst together with oxalate metabolism. This fits with degradation of lignin (and humic compounds) before and during bursts, where ligninolytic action of a synchronized mycelium increases accessibility of carbohydrates together with pectinase and cutinase genes. Recent insights in the decay strategy of brown rot fungi points to the production of hydroxyl radicals by their hyphal front while the bulk of (hemi-) cellulases is produced by older mycelium (Zhang *et al*., 2016, 2019; Presley and Schilling, 2017; Castaño *et al*., 2018). H_2_O_2_ and other radical species can damage proteins involved in polysaccharide deconstruction and spatial separation of radical generation and (hemi-) cellulases can protect these CAZYs from oxidative damage (Zhang *et al*., 2016; Castaño *et al*., 2018). In addition, the pre-treatment of plant cell walls with Fenton chemistry derived radicals or a ligninolytic system allows a more efficient (hemi-) cellulose deconstruction by CAZYs. Therefore, temporal cycling in ligninolytic and cellulolytic activities by *A. bisporus* may enable the repeated pre-treatment of compost and protect extracellular proteins from radical species, resulting in a more efficient substrate decomposition when compared to a spatial separation of ligninolytic and cellulolytic activities.

## Acknowledgements

This research was supported by the Dutch Technology Foundation STW (Grant 11108), which is part of the Netherlands Organization for Scientific Research (NWO), and which is partly funded by the Ministry of Economic Affairs.

## Author Contributions

Conceptualization: AMV, LGL, HABW. Methodology: AMV, RJB, KCH, HS, LGL. Investigation: AMV, RJB, KCH, KS. Formal Analysis: AMV, RAO. Writing: AMV, RJB, KCH, RAO, KS, HABW. Visualization: AMV, RJB, KCH, KS. Funding acquisition: HABW.

