## Supplemental material for "Cycling in degradation of organic polymers and uptake of nutrients by a litter-degrading fungus"

**Supplementary text 1:** Identification of CAZYs and metabolic genes.

Putative activities of previously identified *A. bisporus* CAZYs and proteins involved in carbon metabolism (Patyshakuliyeva *et al.*, 2013, 2015) were inferred by blasting characterized proteins from the CAZY database (29; http://www.cazy.org) to the *A. bisporus* genome and the UniProtKB/Swiss-Prot database (Dataset S2, S8; (Bateman *et al.*, 2017)). The annotation of the functional class of metabolic genes was improved by the identification of an additional D-galacturonate reductase (Protein ID 194338) that was found as the best bi-directional hit of the characterized D-galacturonate reductase (EC 1.1.1.365) of *Hypocrea jecorina* (Kuorelahti *et al.*, 2005). This protein is also a putative DL-glyceraldehyde reductase (EC 1.1.1.372). Formate dehydrogenase (Protein ID 150620) was identified by blastp using the FDH1 gene of *Candida boidinii* (Sakai *et al.*, 1997); protein ID 239344 as improved gene model. Glyoxalases were identified using Pfam 00903. The best bi-directional hit of glyoxylate reductase (EC 1.1.1.26; Protein ID 151018) was identified using GOR1 of *S. cerevisiae,* while glycolaldehyde dehydrogenase could not be identified due to close similarity of aldehyde dehydrogenases. Delta 9-desaturase (Protein ID 194591) and delta 12-desaturase (Protein ID 193825) of *A. bisporus* were identified using the delta 9-desaturase OLE1 of *S. cerevisiae* and the delta 12-desaturase from *Mortierella alpina* (Sakuradani *et al.*, 1999), respectively.

**Supplementary text 2:** Synchronized cell cycle in compost

Genes with GO terms related to cyclins were overrepresented in genes upregulated after a burst (Table S4). Cyclins are involved in regulation of the cell cycle and growth. The role of 14 genes with a cyclin Pfam domain was inferred by blasting the proteins using the UniProtKB/Swiss-Prot database. This identified 5 out of 14 differentially expressed genes as cyclins including 3 group I cyclins that are key cell cycle regulators (Paolillo et al., 2018). By blasting known cell cycle proteins from Aspergillus nidulans (Dörter and Momany, 2016) to the genome of A. bisporus and identifying best bi-directional hits, 4 additional genes related to the cell cycle were identified. This increased the total set of differentially expressed, cell cycle related genes to 8. These included 5 genes that were downregulated 2.6-9.3 fold and 3 genes that were upregulated 3.4-690 fold (Table S9).

**Supplementary text 3:** Compost energetics

Heat production can be related to O_2_ consumption and vice versa using Thorntons Rule of 455 ±15 kJ released heat per mol O_2_ consumed (Thornton, 1917; Hansen et al., 2004). Background subtracted O_2_ consumption measured with the respirometer was used to calculate heat production during a burst (Table S1). To this end, the rate of O_2_ consumption before and after a burst were averaged and subtracted from the O_2_ consumption during bursts to obtain the additional O_2_ consumption during a burst. The sum of additional O_2_ consumption during bursts was 7.9 % of total consumed O_2_. In addition, the temperature increase in compost during a burst was used to calculate additional O_2_ consumption during a burst (Table S2). The maximal consumption of 12.69 mmol O_2_ kg^-1^ compost for a burst that was calculated for compost is close to the consumption of 12.96 – 17.08 mmol O_2_ kg^-1^ compost measured using the largest burst in the respirometer. O_2_ consumption calculated for bursts based on heat release was lower compared to the O_2_ consumption measured by the respirometer. This is expected as heat loss from the uninsulated boxes to the environment by evaporation or otherwise cannot be accounted for. Therefore, O_2_ consumption can fully explain heat production during respiratory bursts.

The RQ of the additionally released CO_2_ and consumed O_2_ was higher than that of the background RQ. At this point it is not possible to identify the substrates being utilized before, during, and after bursts. However, the increased RQ (before or after background subtraction) together with the increase in O_2_ consumption, and CO_2_ and heat release shows that there is increased consumption of more oxidized substrates during bursts as compared to the period in between bursts.

Using the mass balance of (Jurak et al., 2015) and expected heat production of lignin and carbohydrates (Table S3), the heat released from lignin, cellulose, and hemicellulose degradation during a typical PIII was estimated (Table S4). Loss of lignin contributes up to 5 times more heat as compared to cellulose and hemicellulose combined (using klason lignin as estimate for lignin loss). Since the O_2_ consumption during bursts comprised 21 % of total O_2_ consumption (7.9 % of all O_2_ was consumed above the background O_2_ consumption during bursts; Table S1) lignin cannot only be degraded during bursts.

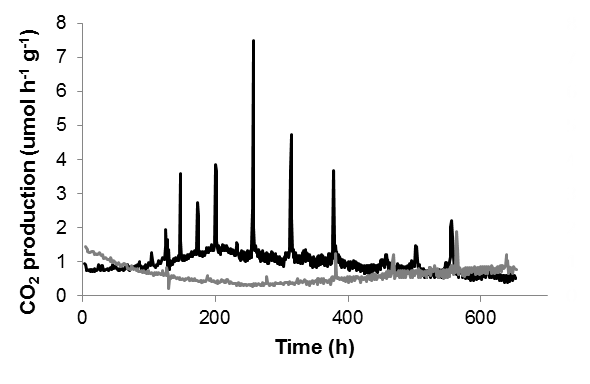

**Figure S1.** CO_2_ production rate of 10 g PII-end compost that had (grey line) or had not (black line) been sterilized prior to inoculation with 4 spawn grains of *A. bisporus*. CO_2_ production was monitored after pre-growth at 25 ˚C for 3 days.

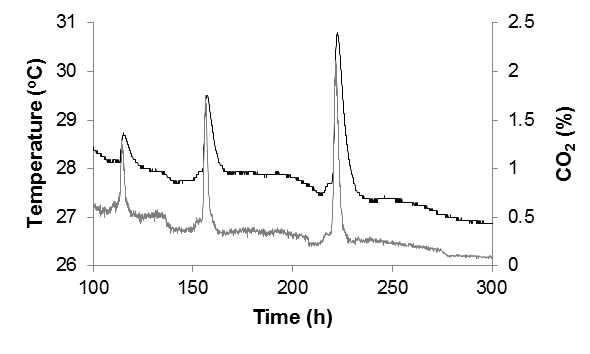

**Figure S2.** Respiratory bursts in 1 kg compost as measured by monitoring changes in temperature (black) and CO_2_ (grey).

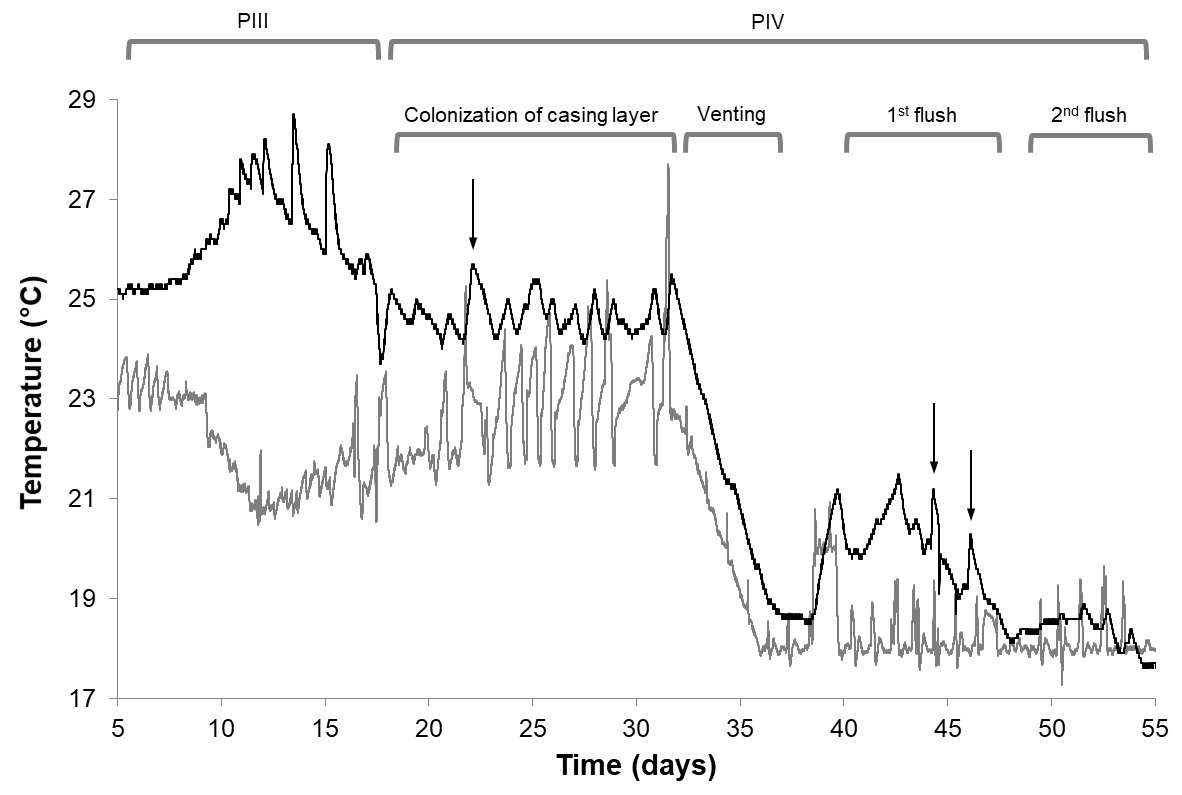

**Figure S3.** Temperature profile of compost (black line) and air in the incubation room (grey line) during vegetative growth (PIII and colonization of casing layer) and mushroom production (Venting, 1^st^ flush, and 2^nd^ flush) of *A. bisporus*. Respiratory bursts occurring in PIV are indicated with an arrow. These temperature peaks cannot be explained by peaks in the temperature of the incubation room.

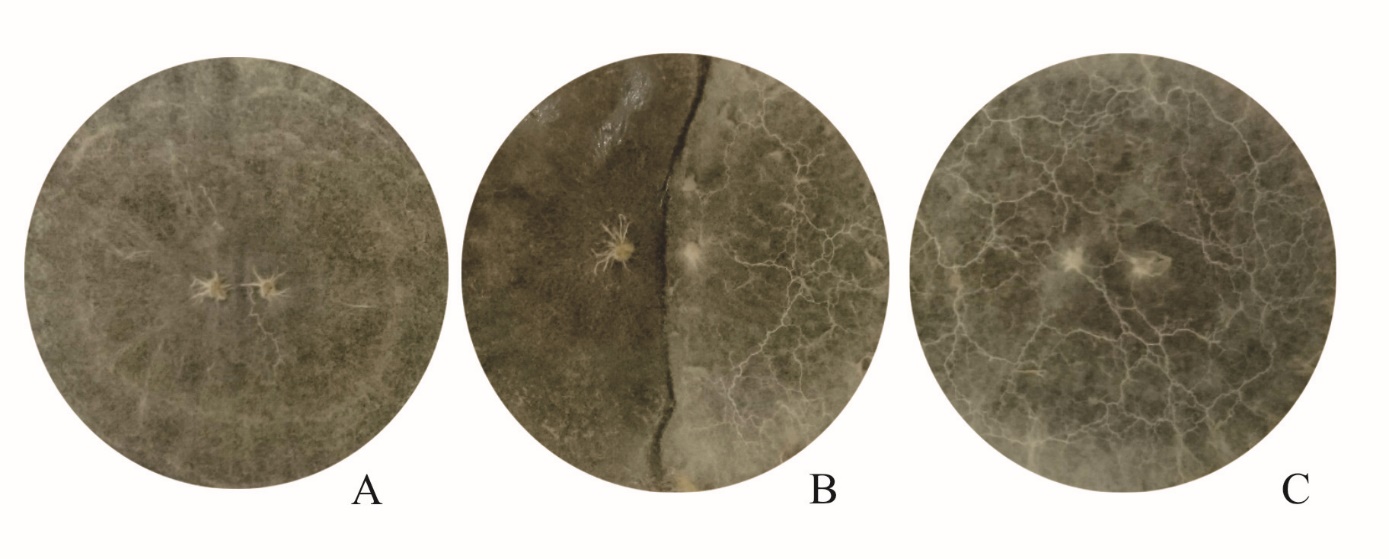

**Figure S4.** Examples of a compatible interactions of strain A15 (A), incompatible interaction of strains A15 and Bisp015 (B) and compatible interaction of strain Bisp015 (C).

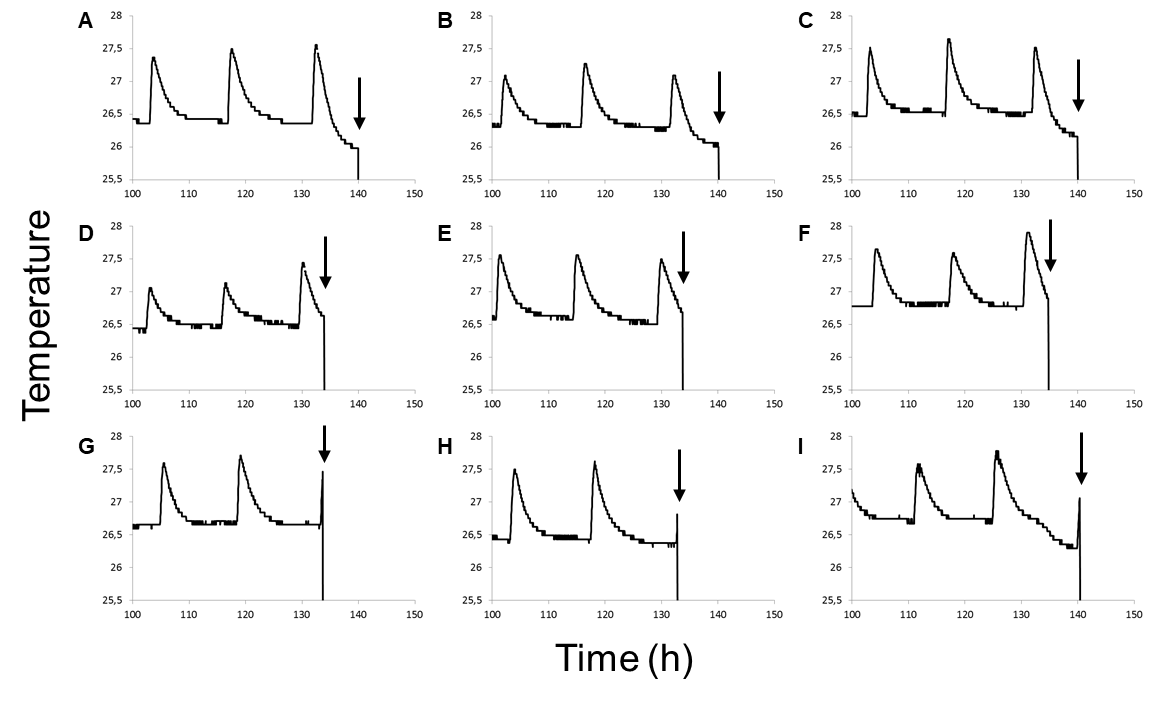

**Figure S5.** Temperature profiles of PIII compost used for RNA isolation in triplicate of inter-burst (A-C), after-burst (D-F), and burst (G-I) stages. Harvesting was done at time points indicated with an arrow. Temperature measurements were started 4 days after inoculation.

**Table S1:** Additionally released CO_2_ (Add. CO_2_) and O_2_ (Add. O_2_) during bursts observed in the respirometer (Figure 1). kJ released per burst was calculated assuming 455 kJ mol^-1^ O_2_ (Hansen *et al.*, 2004). Total CO_2_ production and O_2_ consumption was 831 and 1205 mmol kg^-1^ compost, respectively. The total O_2_ production during bursts was 255.15 mmol kg^-1^ compost and the total of additional O_2_ released during bursts was 95.65 mmol kg^-1^ compost.

| **Burst** | **Total mmol O_2_ consumption kg^-1^ compost per burst** | **Additional CO_2_ mmol kg^-1^ compost per burst** | **Additional O_2_ mmol consumption kg^-1^ compost per burst** | **RQ*** | **Calculated heat release in kJ kg^-1^ compost per burst** |
| --- | --- | --- | --- | --- | --- |
| **33 h** | 9.63 | 0.85 | 0.83 | 1.03 | 0.38 |
| **50 h** | 10.70 | 1.22 | 1.65 | 0.74 | 0.75 |
| **68 h** | 12.41 | 1.59 | 2.10 | 0.76 | 0.96 |
| **88 h** | 8.71 | 2.07 | 2.84 | 0.73 | 1.29 |
| **108 h** | 11.00 | 2.99 | 4.56 | 0.65 | 2.08 |
| **128 h** | 20.53 | 4.30 | 5.08 | 0.84 | 2.31 |
| **148 h** | 19.52 | 7.78 | 10.45 | 0.74 | 4.76 |
| **170 h** | 31.68 | 9.40 | 10.85 | 0.87 | 4.94 |
| **191 h** | 32.29 | 11.96 | 12.96 | 0.92 | 5.90 |
| **211 h** | 27.26 | 7.56 | 8.85 | 0.85 | 4.03 |
| **236 h** | 31.83 | 13.82 | 15.13 | 0.91 | 6.88 |
| **288 h** | 31.09 | 14.50 | 17.08 | 0.85 | 7.77 |
| **380 h** | 8.55 | 2.49 | 3.27 | 0.76 | 1.49 |

*respiratory quotient of the additionally released CO_2_ and consumed O_2_.

**Table S2:** Average (and standard deviation) of the duration of temperature increase, temperature increase, kJ released per burst assuming 60 % water content, and mmol O_2_ kg^-1^ compost released per burst. Values represent the average of the 6 sensors in Figure 2 for a burst around a time point. The maximum values for one of the peaks at 218 h are shown.

| **Burst** | **Period of T* increase (min)** | **dT**** | **Heat release in kJ kg^-1^ compost per burst** | **mmol O_2_ kg^-1^ compost per burst** |
| --- | --- | --- | --- | --- |
| **73 h** | 49.3 (6.3) | 0.81 (0.06) | 2.02 (0.15) | 4.44 (0.34) |
| **88 h** | 51.9 (5.6) | 0.83 (0.12) | 2.06 (0.31) | 4.53 (0.67) |
| **103 h** | 48.9 (6.6) | 0.89 (0.14) | 2.22 (0.36) | 4.87 (0.79) |
| **120 h** | 41.1 (6) | 0.44 (0.07) | 1.11 (0.18) | 2.44 (0.39) |
| **137 h** | 59.4 (3.1) | 1.51 (0.05) | 3.78 (0.12) | 8.3 (0.26) |
| **156 h** | 46.7 (11.3) | 0.23 (0.14) | 0.57 (0.34) | 1.25 (0.75) |
| **175 h** | 69.8 (7.7) | 1.39 (0.26) | 3.47 (0.64) | 7.63 (1.42) |
| **218 h** | 85.5 (14.5) | 1.13 (0.7) | 2.82 (1.76) | 6.2 (3.86) |
| **Maximum (218 h)** | 94.1 | 2.31 | 5.78 | 12.69 |

*temperature in °C; **increase in compost temperature in °C.

**Table S3:** Estimates of lignin, cellulose, and hemicellulose heat production upon mineralization. Numbers are based on data from (Ioelovich, 2018) and Thorntons Rule (Thornton, 1917).

| **Monomer** | **Monomer formula** | **Monomer MW** | **Mol O_2_ per mol mineralized monomer** | **Heat production in kJ mol^-1^ based on** (Ioelovich, 2018) | **Heat production in kJ mol^-1^ based on Thorntons Rule** |
| --- | --- | --- | --- | --- | --- |
| Lignin | C_10.24_H_12.48_O_3.24_ | 188.29 | 11.74 | -5452 | -5342 |
| Cellulose | C_6_H_10_O_5_ | 162 | 6 | -2988 | -2730 |
| Hemicellulose | C_5_H_8_O_4_ | 132 | 5 | -2477 | -2275 |

**Table S4:** Estimated heat production as a consequence of lignin, cellulose, and hemicellulose loss during a typical PIII. Values for the PIII mass balance were taken from (Jurak *et al.*, 2015). Conversion to heat production are based on Table S3.

| **Component** | **PII mass balance (g)** | **PIII mass balance (g)** | **Loss relative to PII (g)** | **% loss during PIII** | **mol "monomer"** | **kJ heat kg^-1^ compost*** | **O_2_****  **mol kg^-1^** |
| --- | --- | --- | --- | --- | --- | --- | --- |
| Total (with Klason lignin) | 787 | 708 | 79 | 10.0% |  | -1305 | -2.9 |
| Lignin (pyrogram based) | 208 | 91 | 117 | 56.3% | 0.62 | -3387 | -7.4 |
| Lignin (klason based) | 252 | 214 | 38 | 15.1% | 0.20 | -1100 | -2.4 |
| Cellulose | 197 | 186 | 11 | 5.6% | 0.07 | -114 | -0.2 |
| Hemicellulose | 110 | 104 | 6 | 5.6% | 0.04 | -91 | -0.2 |

*Based on Ioelovich (2018). **Based on Thornton’s rule.

**Dataset S1:** Enrichment analysis of GO and PFAM before, during, and after bursts.

**Dataset S2:** Expression of all CAZYs before, during, and after bursts.

**Dataset S3:** Total number and differentially expressed genes per CAZY class in 4 different

FPKM ranges before, during, and after bursts.

**Dataset S4:** Expression of all genes before, during, and after bursts.

**Dataset S5:** Expression of protease genes before, during, and after bursts.

**Dataset S6:** Expression of beta-etherase (GST), heme-thiolate peroxidase (HTP), and P450

genes before, during, and after bursts.

**Dataset S7:** Expression of transporter genes before, during, and after bursts.

**Dataset S8:** Expression of metabolic genes before, during, and after bursts.

**Dataset S9:** Expression of cell cycle related cyclin genes before, during, and after bursts
